# A comparison of two universal angiosperm bait sets and the phylogenomics of Alismatales

**DOI:** 10.1101/2025.09.15.676180

**Authors:** Ed Biffin, Michelle Waycott, Timothy A. Hammer, Kor-jent van Dijk

## Abstract

High throughput sequencing of hybridisation capture libraries provides an efficient approach for assembling large scale phylogenomic data. These include ‘universal’ bait sets that aim to generate comparable data from any lineage within the taxon of interest. Here, we present the OzBaits v2 bait set, which targets a set of low copy nuclear loci for angiosperms. Using published genomic data, we design a set of RNA baits targeting a single exon in each of 98 putatively orthologous nuclear protein coding genes. We tested the efficiency of this bait set for a diverse range of angiosperms and recovered, on average, 93 (95%) genes per sample. We compared a common set of samples for the monocot order Alismatales enriched using OzBaits and the Angiosperms353 (A353) bait set, a widely used universal probe set targeting up to 353 nuclear genes in angiosperms. Gene recovery was, on average, c. 1.7 times higher for OzBaits relative to A353. Using proxies for signal and bias to rank gene alignments by their phylogenetic usefulness, we found that on average, the OzBaits data had higher phylogenetic utility. Both data sets resolved largely congruent, well-supported phylogenies for Alismatales although measures of internal discordance where higher for the A353 data. We discuss the implications of these findings for the design universal baits sets.

## Introduction

With the advent of efficient sequencing technologies and the development of large-scale (genomic) data sets, reduced representation sequencing approaches have become a mainstay of evolutionary biology research (McKain et al. 2018). Reduced representation methods including transcriptomics, restriction-site associated DNA sequencing (e.g. RAD-seq), targeted amplicon sequencing and hybridisation-capture, allow the sequencing of a reduced portion of the genome across many samples for the equivalent effort of a single whole genome. Hybridisation-capture experiments target genetic loci of interest, such as phylogenetically informative low-copy nuclear genes (LCNG), using specifically designed RNA bait sequences that hybridise in solution with targeted regions and are enriched prior to high-throughput sequencing (Weitemier et al. 2014; Johnson et al. 2019). Hybridisation bait sets for evolutionary studies include those that target major linages spanning many millions of years (so-called ‘universal’ bait sets, e.g. Buddenhagen et al. 2016; Wolf et al. 2018; Johnson et al. 2019; Liu et al. 2019; Brienholt et al. 2021; Hutter et al. 2021; Khan et al. 2024) through to bait sets that target lower taxa ranging from family (e.g. Choi et al. 2020; Fonseca et al. 2023), generic (e.g. Michel et al. 2022) and species-specific approaches (e.g. Hill et al. 2019). On the one hand, a universal bait set facilitates the recovery of phylogenetically useful data from divergent evolutionary lineages enabling the development of large-scale comparable sequence data sets broadly across the taxon of interest (e.g. Zuntini et al. 2024). On the other hand, bait sets with a narrower taxonomic focus require fewer baits per gene of interest and tend to show higher enrichment efficiency (i.e. the number of ‘on target’ reads as a proportion of the total) because of lower divergence between bait and target sequences (Liu et al. 2019).

Several studies to date have compared the relative performance of ‘universal’ versus ‘taxon specific’ bait sets (e.g. Larridon et al. 2020; Shah et al. 2021; Yardeni et al. 2022) although few have contrasted the performance of ‘universal’ bait sets that employ different strategies for bait design. A key issue for bait set design is efficiency in terms of cost per base pair of sequence data generated and this is a function of the number of loci that are targeted and the level of genetic divergence between the bait and target sequences. Thus, for a given unit cost, increasing the number of target loci (or number of base pairs targeted) can be achieved by reducing the diversity and specificity of probe sequences. Conversely, increasing probe specificity could be achieved by reducing the number of genetic loci included in the bait set. While maximising the amount of data generated is a key consideration in universal bait set design, trade-offs in terms of probe specificity and enrichment efficiency could potentially influence the quality of the data including reduced matrix occupancy, lower base call accuracy and increased proportions of missing data (Hutter et al. 2022). Each of these aspects has potential to impact the phylogenetic utility of the data sets and for instance, fragmentary sequences have been found to increase noise (non-phylogenetic signal) in large phylogenomic data sets (Sayyari et al. 2017).

We present OzBaits v2.0, a universal probe set that is designed to recover c. 100 LCNG from any angiosperm lineage, building on the OzBaits v1.0 bait kit, which targets fewer genes (Waycott et al. 2021). In the present study, we focus on the OzBaits v2.0 nuclear component and we compare the performance of these with a widely used alternative, the Angiosperms353 baits (A353; Johnson et al. 2019), which target up to 353 LCNG in any angiosperm. The former targets a single (or partial) exon per genetic locus and relative to A353 and has higher probe specificity while the latter attempts to recover entire genes by tiling probes for each target locus on transcript sequences. As part of the Genomics for Australian Plants (GAP; https://www.genomicsforaustralianplants.com/) initiative, we have generated sequence data for a broad range of angiosperm species to assess the efficiency of gene recovery using the OzBaits v2.0 probe set. In addition, we compare the performance of OzBaits and A353, focussing on Australian representatives of the monocot lineage Alismatales an in particular, ‘core Alismatales’, a diverse predominantly hydrophytic lineage that includes all known fully submerged marine angiosperms (seagrasses). For each bait set, we have sequenced the same set of samples, employed equivalent library preparation steps and used a common bioinformatics pipeline, enabling meaningful comparisons between OzBaits and A353 in terms of the differences in bait design and how these influence the phylogenetic utility of the data sets generated.

## Methods

### Bait Design and Synthesis OzBaits V2.0

We designed target enrichment probes to cover all angiosperms using published transcriptomes and genomic sequences. Target genes were selected from Duarte et al. (2010) who report a set of putatively orthologous low copy nuclear genes shared in *Arabidopsis, Populus, Vitis* and *Oryza* (APVO SSC genes sensu Duarte et al., 2010). We first examined the corresponding genomic data (coding sequences: CDS) available on Phytozome (*v. 12*, including 48 species of angiosperms encompassing all major flowering plant linages (https://phytozome.jgi.doe.gov) to identify a subset of genes based upon several criteria including: copy number (genes are single copy in the majority of diploid genomes); sequence conservation (c. 70% average pairwise identity) and representation across the sequenced genomes (locus represented in > c. 70% of genomes). For the selected genes, we used the CDS for *Arabidopsis thaliana* (Araport11; Cheng et al. 2017) to retrieve putatively homologous transcript sequences from the 1000 Plants Project (1KP; https://www.onekp.com; Leebens-Mack *et al*. 2019) using the China National Genebank (https://db.cngb.org) BLAST portal and the following settings: Discontiguous Mega-Blast, expect value=10, maximum target sequences=1000, selected organisms=Magnoliophyta (taxid:3398). Additional sequences were sourced from SeagrassDB (Sablok et al. 2018), which includes transcriptome data for marine and aquatic angiosperms from the Alismatales. The sequences retrieved from the above sources were combined with the Phytozome data for each gene and made into a BLAST database in Geneious Prime 2022.0.1 (https://www.geneious.com). We queried each BLAST database using the *A. thaliana* gene family member with exon annotations manually added, and the following settings: Discontiguous Mega-Blast, expect value=10, maximum target sequences=1200, results=Hit Table, retrieve=Matching Region with Annotation. We then extracted sequences matching the exon annotations and selected a single exon per gene with the caveat that exon size was >180 and less than 800 bp, to facilitate probe tiling while reducing the number of required probes per target locus. The extracted sequences were clustered using CD-HIT-EST (Li and Godzik, 2006; http://weizhong-lab.ucsd.edu/cdhit_suite) with a sequence identity cut-off fraction of 0.85 (the approximate limit for genetic divergence between probe and target sequence for efficient hybridisation, Hancock-Hanser et al. 2013) and a length similarity fraction of 0.2, and one representative sequence (the longest) per cluster was selected. A total of 19,652 representative sequences were used for bait design, ranging in length from 180-660 bp, with a mean length of c. 319 bp. Bait design (myBaits) and synthesis was performed by Daicel-Arbor Biosciences (formerly Mycroarray; Ann Arbor, Michigan, USA; Cat. # 300196R.v5) with 100-nucleotide baits and ∼2X flexible tiling density for a total of 101,467 baits (Supplementary material-S1).

### Taxon sampling

We explored the performance of the Ozbaits bait set using a broad sample of 19 angiosperm taxa including monocots (Asphodelaceae: *Caesia, Xanthorrhoea;* Araceae: *Lemna, Typhonium;* Maundiaceae: *Maundia;* Ruppiaceae: *Ruppia*), eudicots (Ceratophyllaceae: *Ceratophyllum*), core eudicots (Dilleniaceae: *Dillenia, Hibbertia*), rosids (Malvaceae: *Androcalva, Lasiopetalum, Thomasia;* Myrtaceae: *Darwinia, Leptospermum*), Caryophyllales (Droseraceae: *Drosera*) and asterids (Plantaginaceae: *Plantago*). We also developed a dataset for Alismatales comprising 96 samples including 11 of the 14 currently accepted families (Stevens, 2001 onwards; http://www.mobot.org/MOBOT/research/APweb/, Butomaceae, Scheuchzeriaceae and Tofieldiaceae are not naturally present in Australia). We included 31 of the 38 genera with Australian representation from the ‘core alismatids’ (Wilson et al. 2011) along with a single representative for each of three aroid genera. *Ceratophyllum* and *Plantago* were included as outgroups (Table 1).

**Table 1:** Details of samples included in this study. The data set ‘both’ includes sequence data generated using the OzBaits and A353 bait sets.

| Taxon | Family | GAP sample id | Voucher specimen | Dataset |
| --- | --- | --- | --- | --- |
| 376498_Aff_Plantago | Scrophulariaceae | 376498 |  | both |
| *Alisma lanceolatum* With. | Alismataceae | 376416 | J.G. Conran 3240 (AD289809) | both |
| *Alisma plantago-aquatica* L. | Alismataceae | 376417 | J.G. Conran 3142 (AD289808) | both |
| *Althenia australis* (J.Drumm. ex Harv.) F.Muell. | Potamogetonaceae | 376418 | GIBS 1975 (AD289811) | both |
| *Althenia bilocularis* (Kirk) Cockayne | Potamogetonaceae | 376419 | GIBS 1976 (AD289802) | both |
| *Althenia cylindrocarpa* (Körn. ex Müll.Berol.) Asch. | Potamogetonaceae | 376420 | GIBS 1947 (AD276764) | both |
| *Althenia marina* (E.L.Robertson) Yu Ito | Potamogetonaceae | 376421 | GIBS 1953 (AD283425) | both |
| *Althenia patentifolia* (E.L.Robertson) T.D.Macfarl. & D.D.Sokoloff | Potamogetonaceae | 376422 | GIBS 1972 (AD289793) | both |
| *Althenia preissii* (Lehm.) F.Muell. | Potamogetonaceae | 376423 | GIBS 1941 (AD276663) | both |
| *Althenia*_sp Conran & Waycott | Potamogetonaceae | 376424 | J.G. Conran 3852 (AD289805) | both |
| *Amphibolis antarctica* (Labill.) Sond. & Asch. | Cymodoceaceae | 376425 | GIBS 1798 (AD264197) | both |
| *Amphibolis griffithii* (J.M.Black) Hartog | Cymodoceaceae | 376426 | GIBS 1803 (AD283592) | both |
| *Androcalva johnsonii* (Guymer) C.F.Wilkins & Whitlock | Malvaceae | 376785 | C.F. Wilkins 2016A (PERTH 07963807) | OzBaits |
| *Aponogeton bullosus* H.Bruggen | Aponogetonaceae | 376427 | R. Booth 2607 (AD132657) | both |
| *Aponogeton distachyos* L.f. | Aponogetonaceae | 376428 | H.J. Eichler 16194 (AD96311389) | both |
| *Aponogeton elongatus* F.Muell. ex Benth. | Aponogetonaceae | 376429 | P.E. Conrick 1108 (AD98322239) | both |
| *Aponogeton tofus* S.W.L.Jacobs | Aponogetonaceae | 376430 | J.G. Conran 3804 (AD289796) | both |
| *Blyxa aubertii* Rich. | Hydrocharitaceae | 376431 | D.E. Symon 7921 (AD98666314) | both |
| *Blyxa japonica* (Miq.) Maxim. ex Asch. & Gürke | Hydrocharitaceae | 376433 | J.G. Conran 3798 (AD289812) | both |
| *Blyxa octandra* (Roxb.) Thwaites | Hydrocharitaceae | 376432 | C.L. Dunlop & L. Craven 5898 (AD98231145) | both |
| *Caesia setifera* Baker | Asphodelaceae | 377222 | B.S. Wannan 5615 (BRI AQ0913591) | OzBaits |
| *Caldesia oligococca* (F.Muell.) Buchenau | Alismataceae | 376434 | T.A. Halliday 473 (AD98232154) | both |
| *Ceratophyllum demersum* L. | Ceratophyllaceae | 376508 | J.G. Conran 3211 (AD289814) | both |
| *Cycnogeton alcockiae* (Aston) Mering & Kadereit | Juncaginaceae | 376435 | D.E. Murphet 4507(AD156411) | both |
| *Cycnogeton dubium* (R.Br.) Mering & Kadereit | Juncaginaceae | 376436 | H.I. Aston 2745 (AD99411237) | both |
| *Cycnogeton huegelii* Endl. | Juncaginaceae | 376437 | T.A. Halliday 176 (AD97611053) | both |
| *Cycnogeton lineare* (Endl.) Sond. | Juncaginaceae | 376438 | A.E. Orchard 1417 (AD97111020) | both |
| *Cycnogeton microtuberosum* (Aston) Mering & Kadereit | Juncaginaceae | 376439 | H.I. Aston 2815 (AD99411241) | both |
| *Cycnogeton multifructum* (Aston) Mering & Kadereit | Juncaginaceae | 376440 | E.M. Canning 6388 (AD98950269) | both |
| *Cycnogeton procerum* (R.Br.) Buchenau | Juncaginaceae | 376441 | J.G. Conran 3812 (AD289800) | both |
| *Cycnogeton rheophilum* (Aston) Mering & Kadereit | Juncaginaceae | 376442 | H.J. Eichler 13316 (AD95917002) | both |
| *Cymodocea rotundata* Ehrenb. & Hempr. ex Asch. & Schweinf. | Cymodoceaceae | 376443 | K-j van Dijk s.n. (AD272801) | both |
| *Cymodocea serrulata* (R.Br.) Asch. & Magnus | Cymodoceaceae | 376444 | GIBS 1349 (AD272845) | both |
| *Damasonium minus* (R.Br.) Buchenau | Alismataceae | 376445 | J.G. Conran 3936 (AD289820) | both |
| *Darwinia diosmoides* (DC.) Benth. | Myrtaceae | 376930 | E. Massenbauer 763 (PERTH 9171177) | OzBaits |
| *Dillenia alata* (R.Br. ex DC.) Martelli | Dilleniaceae | 376560 | J.O. Westaway 3132 (DNA D0195012) | OzBaits |
| *Drosera binata* Labill. | Droseraceae | 376553 | F.J. Nge 1402 (AD286718) | OzBaits |
| *Drosera scorpioides* Planch. | Droseraceae | 376531 | J.G. Conran 4159 (AD 286723) | OzBaits |
| *Egeria densa* Planch. | Hydrocharitaceae | 376446 | J.G. Conran 3796 (AD289810) | both |
| *Enhalus acoroides* (L.f.) Royle | Hydrocharitaceae | 376447 | GIBS 492 (AD272358) | both |
| *Halodule uninervis* (Forssk.) Asch. | Cymodoceaceae | 376448 | GIBS 505 (AD284671) | both |
| *Halophila australis* Doty & B.C.Stone | Hydrocharitaceae | 376449 | GIBS 1821 (AD283603) | both |
| *Halophila capricorni* Larkum | Hydrocharitaceae | 376450 | GIBS 36 (AD272471) | both |
| *Halophila decipiens* Ostenf. | Hydrocharitaceae | 376451 | GIBS 1052 (AD269769) | both |
| *Halophila decipiens* | Hydrocharitaceae | 376454 | GIBS 1776 (AD289788) | both |
| *Halophila minor* (Zoll.) Hartog | Hydrocharitaceae | 376452 | GIBS 958 (AD272420) | both |
| *Halophila ovalis* (R.Br.) Hook.f. | Hydrocharitaceae | 376453 | GIBS 1829 (AD283389) | both |
| *Halophila spinulosa* (R.Br.) Asch. | Hydrocharitaceae | 376455 | GIBS 47 (AD272467) | both |
| *Halophila tricostata* Greenway | Hydrocharitaceae | 376456 | GIBS 34 (AD272472) | both |
| *Heterozostera nigricaulis* J.Kuo | Zosteraceae | 376457 | V. Stajsic 4264 (AD217053) | both |
| *Heterozostera polychlamys* J.Kuo | Zosteraceae | 376458 | GIBS 1955 (AD283408) | both |
| *Hibbertia crinita* Toelken | Dilleniaceae | 376579 | T.A. Hammer, A.E. McDougal 194 (AD286612) | OzBaits |
| *Hibbertia salicifolia* (DC.) F.Muell. | Dilleniaceae | 376636 | P.I. Forster 30223 (AD196618) | OzBaits |
| *Hydrilla verticillata* (L.f.) Royle | Hydrocharitaceae | 376459 | J.G. Conran 3792 (AD289801) | both |
| *Hydrocleys nymphoides* (Humb. & Bonpl. ex Willd.) Buchenau | Alismataceae | 376460 | J.G. Conran 3801 (AD289819) | both |
| *Lasiopetalum pterocarpum* E.M.Benn. & K.A.Sheph. | Malvaceae | 376720 | C. Wilkins 2108 (PERTH 07220219) | OzBaits |
| *Lemna disperma* Hegelm. | Araceae | 376510 | C. Austin s.n. (AD281877) | both |
| *Leptospermum polygalifolium* subsp. *tropicum* Joy Thomps. | Myrtaceae | 376911 | B.S. Wannan 6676 (CNS 143594.1) | OzBaits |
| *Limnobia laevigatum* (Humb. & Bonpl. ex Willd.) Heine | Hydrocharitaceae | 376461 | J.G. Conran 4429 (AD289797) | both |
| *Maundia triglochinosides* F.Muell. | Maundiaceae | 376462 | GIBS 1974 (AD289791) | both |
| *Najas tenuifolia* R.Br. | Hydrocharitaceae | 376463 | J.G. Conran 3793 (AD289799) | both |
| *Ottelia alismoides* (L.) Pers. | Hydrocharitaceae | 376464 | C.R. Dunlop & L. Craven 5937 (AD98231141) | both |
| *Ottelia ovalifolia* (R.Br.) Rich. | Hydrocharitaceae | 376465 | J.G. Conran 3840 (AD289806) | both |
| *Posidonia angustifolia* Cambridge & J.Kuo | Posidoniaceae | 376466 | GIBS 1831 (AD283898) | both |
| *Posidonia australis* Hook.f. | Posidoniaceae | 376467 | GIBS 1003 (AD272314) | both |
| *Posidonia coriacea* Cambridge & J.Kuo | Posidoniaceae | 376468 | GIBS 1367 (AD272280) | both |
| *Posidonia denhartogii* J.Kuo & Cambridge | Posidoniaceae | 376469 | GIBS 1833 (AD284660) | both |
| *Posidonia ostenfeldii* Hartog | Posidoniaceae | 376470 | GIBS 1834 (AD283897) | both |
| *Posidonia robertsoniae* J.Kuo & Cambridge | Posidoniaceae | 376471 | GIBS 1441 (AD272581) | both |
| *Posidonia sinuosa* Cambridge & J.Kuo | Posidoniaceae | 376472 | GIBS 1845 (AD283404) | both |
| *Potamogeton australiensis* A.Benn. | Potamogetonaceae | 376473 | W.M. Curtis s.n. (AD99537017) | both |
| *Potamogeton crispus* L. | Potamogetonaceae | 376474 | C.R. Alcock 10006 (AD98916229) | both |
| *Potamogeton drummondii* Benth. | Potamogetonaceae | 376475 | M. Angeloni s.n. (AD206122) | both |
| *Potamogeton ochreatus* Raoul | Potamogetonaceae | 376476 | R. Taylor 531 (AD110788) | both |
| *Potamogeton octandrus* Poir. | Potamogetonaceae | 376477 | M. Fagg 733 (AD97051304) | both |
| *Potamogeton perfoliatus* L. | Potamogetonaceae | 376478 | AC Beaglehole & T Henshall 33426 (AD98009127) | both |
| *Potamogeton reduncus* Hagstr. | Potamogetonaceae | 376479 | C.R. Alcock 2817 (AD97007024) | both |
| *Potamogeton sulcatus* A.Benn. | Potamogetonaceae | 376480 | S.E. Papassotiriou & S.W.L. Jacobs 105 (AD255359) | both |
| *Potamogeton tepperi* A.Benn. | Potamogetonaceae | 376481 | Mr Barbi s.n. (AD160168) | both |
| *Potamogeton tricarlinatus* F.Muell. & A.Benn. | Potamogetonaceae | 376482 | D.E. Murphet, R.L. Taplin 3808 (AD112069) | both |
| *Ruppia maritima* L. | Ruppiaceae | 376483 | GIBS 1878 (AD276770) | both |
| *Ruppia megacarpa* R.Mason | Ruppiaceae | 376484 | GIBS 1932 (AD289756) | both |
| *Ruppia polycarpa* R.Mason | Ruppiaceae | 376485 | GIBS 1933 (AD289757) | both |
| *Ruppia* sp. Waycott & Conran | Ruppiaceae | 376486 | GIBS 1973 (AD289792) | both |
| *Ruppia tuberosa* J.S.Davis & Toml. | Ruppiaceae | 376487 | E. O'Loughlin, R. Lewis 9505 (AD289815) | both |
| *Sagittaria platyphylla* (Engelm.) J.G.Sm. | Alismataceae | 376488 | J.G. Conran 4388 (AD289813) | both |
| *Stuckenia pectinata* (L.) Borner | Potamogetonaceae | 376489 | S. Clarke s.n. (AD273818) | both |
| *Syringodium isoetifolium* (Asch.) Dandy | Cymodoceaceae | 376490 | GIBS 10 (AD272803) | both |
| *Thalassia hemprichii* (Ehrenb.) Asch. | Hydrocharitaceae | 376491 | GIBS 24 (AD268034) | both |
| *Thalassodendron ciliatum* (Forssk.) Hartog | Cymodoceaceae | 376492 | GIBS 500 (AD272366) | both |
| *Thalassodendron pachyrhizum* Hartog | Cymodoceaceae | 376493 | GIBS 90 (AD272218) | both |
| *Thomasia solanacea* (Sims) J.Gay | Malvaceae | 376771 | K.A. Shepherd 1872 (PERTH 09427953) | both |
| *Triglochin hexagona* J.M.Black | Juncaginaceae | 376494 | R.J. Bates 47523 (AD99803327) | both |
| *Triglochin isingiana* (J.M.Black) Aston | Juncaginaceae | 376495 | D.E. Murphet 6906 (AD238756) | both |
| *Triglochin longicarpa* (Ostenf.) Aston | Juncaginaceae | 376496 | P. Wilson 3542 (AD96525160) | both |
| *Triglochin minutissima* F.Muell. | Juncaginaceae | 376497 | J.G. Conran 3228 (AD289818) | both |
| *Triglochin nana* F.Muell. | Juncaginaceae | 376499 | BS23-28392 (AD113217) | both |
| *Triglochin stowardii* N.E.Br. | Juncaginaceae | 376500 | E.A. Shaw 554 (AD99212071) | both |
| *Triglochin trichophora* Nees | Juncaginaceae | 376501 | D.E. Murphet 1279 (AD99151121) | both |
| *Triglochin turrifera* Ewart | Juncaginaceae | 376502 | D.E. Murphet, R.L. Taplin 2061 (AD99605142) | both |
| *Typhonium alismifolium* F.Muell. | Araceae | 376511 | BSOP-314 (AD111184) | both |
| *Typhonium cochleare* A.Hay | Araceae | 377245 | N.J. Cuff 1423 (DNA D0273539) | both |
| *Typhonium taylorii* A.Hay | Araceae | 377268 | K. Brennan 11992 (DNA D0285606) | both |
| *Vallisneria australis* S.W.L.Jacobs & Les | Hydrocharitaceae | 376503 | D.E. Murphet 3534 (AD107503) | both |
| *Vallisneria caulescens* F.M.Bailey & F.Muell. ex F.M.Bailey | Hydrocharitaceae | 376504 | R.M. Barker 442 (AD98532225) | both |
| *Vallisneria nana* R.Br. | Hydrocharitaceae | 376506 | R. Taylor (AD289755) | both |
| *Xanthorrhoea preissii* Endl. | Asphodelaceae | 377209 | T. McLay 086 (MELUM125730a) | OzBaits |
| *Zannichellia palustris* L. | Potamogetonaceae | 376505 | J.G. Conran 3794 (AD289798) | both |
| *Zantedeschia aethiopica* (L.) Spreng. | Araceae | 376509 | C.J. Brodie, J. Walter 7961 (AD279096) | both |
| *Zostera muelleri* Irmisch ex Asch. | Zosteraceae | 376507 | E.L. Robertson s.n. (AD99023051) | both |

### DNA extraction, library preparation, and sequencing

DNA extractions and library preparations were done at two separate institutions. Initial OzBaits performance testing and non-GAP sample processing was done at the University of Adelaide ADIFF (Advanced DNA, Identification and Forensic Facility) and the GAP initiative samples were processed at the Australian Genome Research Facility (AGRF, Melbourne, Australia) as part of Bioplatforms Australia (Sydney, Australia). At the ADIFF DNA was extracted using the DNeasy® Plant mini kit (Qiagen) as per the manufacturer’s instructions using 20-30 mg of dry tissue. Samples were ground in 2.0 mL screw cap tubes using a Bead Ruptor 24 (Omni International, Kennesaw Georgia, USA) with zirconia beads. DNAs where then converted into genomic libraries using a customized and miniaturized library preparation process that was developed at the University of Adelaide for cost reduction. Detailed laboratory protocols and sequencing adapter designs can be found at DOI: dx.doi.org/10.17504/protocols.io.dm6gpbzm8lzp/v1 (Private link for reviewers: https://www.protocols.io/private/1FB56482A98611EC922E0A58A9FEAC02 to be removed before publication.). In brief, library preparations were done with the NEBNext® Ultra™ II FS DNA Library Prep kit with Fragmentase and Sample Purification Beads (New England Biolabs, Ipswich, MA, USA). Neat DNA extracts were used as the starting material and reactions were done in 1/3 volumes. To enable bioinformatics processing following hybrid capture, custom stubby Y-adaptors with synthetic “barcodes” were annealed to the ends of the DNA fragments and amplified to create half-completed libraries (index and p5 and p7 grafting sites missing). Samples were pooled 16-plex and hybrid capture performed with the myBaits custom OzBaits_NR set (Cat. # 300496R.V5) probes following myBaits manufactures manual with V5 chemistry. Hybridization was done at 65°C and incubated for 24 h. Post capture PCR was performed on the half build libraries by fusing the i5 and i7 indexes and grafting sites to the ends of the DNA fragments. Libraries were pooled in equimolar concentrations and size selected to 350–600 bp, quantified on a 2100 Bioanalyzer (Agilent) and sent for sequencing.

For the GAP initiative samples dried plant tissue (20–30 mg) was provided to AGRF and ground using a TissueLyser II (Qiagen) with tungsten carbide beads. Genomic DNA was extracted using the DNeasy® Plant mini kit (Qiagen) as per the manufacturer’s instructions on a QIAcube Connect (Qiagen). DNA quantity and quality was assessed using 1% E-gel with Sybr Safe dye (Thermo Fisher) and concentrations assessed using Quantifluor dsDNA assay (Promega). Libraries were prepared using the NEBNext Ultra II FS Library Prep Kit (New England Biolabs, Ipswich, MA, USA), following the manufacturer’s instructions targeting inserts of approximately 350 bp. The libraries were enriched using the myBaits custom OzBaits_NR (Cat. # 300496R.v5) and myBaits Expert Plant Angiosperms353 v1 (Cat. # 308108.v5) (Johnson et al. 2019) bait sets with V5 chemistry. Pooled libraries (12–16 plex) were enriched by hybridising at 65°C (A353) and 64°C (Ozbaits_NR) respectively. All sequencing (ADIFF and AGRF) was done on a NovaSeq 6000 (Illumina Inc., San Diego, USA) with v1.5 chemistry and 150 bp paired-end reads.

### Bionformatics processing

High-throughput 150 bp paired-end reads were imported into CLC Genomics Workbench v20.0.2 (https://digitalinsights.qiagen.com/) for demultiplexing and trimming using a quality score limit of 0.05 (Phred score c. 13). Reads for each individual were randomly sampled to 4 million reads. The Captus pipeline (v1.0.1; Oritz et al. 2023) was used to assemble the cleaned sequence reads, then extract and align the target regions. Relative to other commonly used tools for assembling hybrid-capture data sets (e.g. SECAPR, Andermann et al. 2018; HybPiper, Johnson et al. 2016), Captus performs well across a range of data types (Oritz et al. 2023), making it valuable in comparing bait sets that use different design strategies.

Reads for each individual were *de novo* assembled using Megahit (v1.2.9; Li *et al*. 2015) (CAPTUS ‘assemble’ function) with the CAPSKIM default *–k-list* (https://edgardomortiz.github.io/captus.docs/assembly/assemble/) with both –*min-count* and *– prune-level* both set to 3. For the extraction step, we used *--nuc_min_identity* 70 and *--nuc_min_score* 2.0 to match the targeted nuclear regions with Scipio (v1.4; Keller et al. 2008). The target file for the A353 was sourced from the Kew Tree of Life Explorer (Baker et al. 2022) and comprised sequences for core Alismatales including representatives of Alismataceae, Aponogetonaceae, Cymodoceaceae, Hybrocharitaceae, Juncaginaceae, Maundiaceae, Posidoniaceae, Potomagetonaceae and Zosteraceae (Supplementary material-S2). For completeness, we also compared the recovery of A353 target genes using the Mega353 (McLay et al. 2023) target file, for which a modified version has been included in Captus (Oritiz et al. 2023). To extract the OzBaits data, we used the 18,663 representative sequences used for bait design as the reference file (Supplementary material-S2).

The Captus ‘align’ function was used to generate multiple sequence alignments (MSAs) for the extracted nuclear and plastid gene data using the *-f* (format) flag to generate separate nucleotide MSAs for each target region comprising the coding sequence (exons; NT), the coding region(s) and introns (genes; GE), and GE plus flanking regions (genes flanked; GF). All extracted markers were aligned with MAFFT (Katoh and Stanley 2013) using the *mafft_auto* algorithm. We used the ‘informed’ paralogy filter in Captus, retaining a maximum of 3 paralogs per sample and using ‘*–tolerance* 2.0*’* to remove putative paralogs. Alignments were trimmed with ClipKIT v1.3.0 (Steenwyk *et al*. 2020) using default settings in Captus. Alignments with fewer than 20 sequences were removed from phylogenetic analyses.

As well as comparing locus recovery from the two bait sets, we also estimated average read depth per target locus and per sample. Read depth estimates were generated using the SECAPR *reference_assembly* function (Andermann et al. 2018), which uses the BWA mapper (Li and Durbin, 2010) for reference-based mapping and Picard (broadinstitute.github.io/picard/) for removing duplicate reads. For the mapping step we used the sampled FASTQ reads and activated the *–reference_type sample-specific* flag, which uses the consensus sequence for each sample and locus as output by the Captus pipeline. We used a Mann-Whitney U test, as implemented in the Past software (*v.* 4; Hammer et al. 2001) to test the statistical significance of coverage estimates between bait sets.

### Phylogenomic analysis

For the nuclear data, we first generated a concatenated alignment comprising the three alignment formats output by Captus for the A353 and OzBaits loci. From this, we generated a locus specific tree (hereafter, gene tree) for each partition using IQ-TREE v2.2.3 (Nguyen *et al*. 2015; Chernomor *et al*. 2016) using 1000 ultrafast bootstrap replicates (UFBS; Mihn et al. 2013) to assess branch support. We used the partitioned ‘GE’ alignments to generate a maximum likelihood (ML) species tree estimate using IQ-TREE with the MFP+MERGE flag activated, which seeks to identify the best model for each partition and then merge like partitions (Kalyaanamoorthy et al. 2017). Only the top 10% of partition merging schemes were examined by using the relaxed hierarchical clustering algorithm (--rclust 10; Lanfear *et al*. 2014).

The partitioned alignment, species and gene tree estimates were used as input to *genesortR* (Mongiardino Koch 2021; https://github.com/mongiardino/genesortR), an R script that sorts and subsamples phylogenomic datasets based on properties that quantify phylogenetic usefulness. For nucleotide data, these include 3 proxies for bias (average pairwise patristic distance, saturation, and root-to-tip variance) and 3 proxies for signal (Robinson-Foulds similarity to the IQ-TREE ‘GE’ topology, average bootstrap support, proportion of variable sites) (see Mongiardino Koch 2021, and references therein). From the sorted gene alignments, we generated 3 final datasets for phylogenetic inference: for each of the OzBaits and A353 data, we selected the highest-ranking alignment format for each gene, along with a ‘100-best’ alignment data set comprising the highest-ranking alignment format for the 100 genes top ranked genes, including both OzBaits and A353 targets. We explored the sensitivity of our results to species tree resolution by collapsing poorly supported nodes (UFBS<90) and rerunning *genesortR*. We generated two additional data sets from the original Captus alignments to test for the effect of low-quality data on phylogenetic usefulness. First, we used the Paragone v1.0.0 (https://github.com/chrisjackson-pellicle/ParaGone; Jackson et al. 2023) pipeline, which is based upon the orthology resolution approach developed by Yang and Smith (2014), to refine quality of each of the original alignments output by Captus. Paragone uses HmmCleaner (Di Franco *et al*. 2019), TreeShrink (Mai *et al*. 2018) and TrimAl (Capella-Gutiérrez *et al*. 2009) to remove sequencing and alignment errors and rogue taxa from MSAs. Secondly, we used TreeShrink to remove potentially spurious long-branched terminals from the alignments. For each of these data sets, we generated gene trees using IQ-TREE (as above) and we compared phylogenetic usefulness rankings for each against the original Captus output using *genesortR*.

For each of our final datasets (‘A353-best’, ‘OzBaits-best’ and ‘100-best’), we used IQ-TREE to generate a concatenated ML estimate of the species tree. We first estimated the best fitting partitioned model scheme, as outlined above, and estimated branch support using 1000 UFBS replicates along with the IQ-TREE implementation of the approximate likelihood ratio test (aLRT;; Guindon *et al*. 2010), with interpretation of support following Minh *et al*. (2013) and recommendations in the IQ-TREE manual (http://www.iqtree.org/doc/Frequently-Asked-Questions).

We generated a gene tree for each partition in our datasets using IQ-TREE, with branch support estimated using the approximate Bayes test (aBayes; Anisimova *et al*. 2011). For each data set, the gene trees were used to generate a coalescent-based species tree estimate using Weighted ASTRAL (wASTRAL; Zhang and Mirarab, 2022) as implemented in Accurate Species Tree EstimatoR (ASTER) (https://github.com/chaoszhang/ASTER). wASTRAL uses weighting to reduce the impact of quartets with low support and/or long branches, thereby providing more accurate species tree estimates compared to unweighted inference (Zhang and Mirarab, 2022). We followed the authors recommendations (https://github.com/chaoszhang/ASTER/blob/master/tutorial/astral-hybrid.md) for maximum and minimum values for aBayes supports (1.0 and 0.33, respectively) and used local posterior probability (LPP) to measure internal branch support for the coalescent species tree.

Both the concatenated and coalescent based species tree estimates were further interrogated using quartet sampling (QS; Pease et al. 2018). The QS approach subsamples quartets from the target tree and alignment to produce 4 metrics, Quartet Concordance (QC), Quartet Differential (QD), Quartet Informativeness (QI) and Quartet Fidelity (QF). Taken together, QS scores describe the degree of topological variation in the data (Pease et al. 2018), providing a valuable alternative to measures such as the bootstrap, which may have limited utility among large data sets (Lanfear and Hahn, 2024). We ran QS analyses with 500 replicates and a log-likelihood cut-off of 2.

## Results

### Gene recovery

The OzBaits bait set performed well across a diverse set of angiosperm lineages including monocots and eudicots (Table 1). Average gene recovery ranged from 81 (*Maundia*) to 97 (*Caesia, Plantago,* Dilleniaceae, Malvaceae) with an average of 93 genes (c. 95%) recovered across all samples (Supplementary material-S3).

For the Alismatales OzBaits data, no sequence data were recovered from two samples and less than 20 genes were recovered for each of 3 samples (Supplementary material-S4). This appears to reflect poor sample quality and/or library preparation given that recovery was high for congeners, and these samples also showed poor recovery from the A353 bait set (below). Excluding these, we recovered on average c. 85 (87%) genes per sample across 91 samples with average values per family ranging from 74 (c. 76%) in Hydrocharitaceae to 95 (c. 97%) in Posidoniaceae. By way of comparison, the per sample gene recovery for the A353 data using Alismatales specific references (Supplementary material-S5) averaged 208 (c. 66%) with minimum and maximum average per family values ranging from 147 (47%; Hydrocharitaceae) to 245 (78%; Zosteraceae). Recovery using the mega353 references was substantially lower (Supplementary material-S6) and is not discussed further. In Figure 1 (Supplementary material-S7), we compare proportion of genes recovered per sample from the OzBaits versus the A353 bait sets, by family. The proportion of loci recovered from OzBaits was in all cases higher by a factor of c. 20% (Araceae, Juncaginaceae) and up to c. 80% in Alismataceae, Hydrocharitaceae and Posidoniaceae.

**Figure 1:**
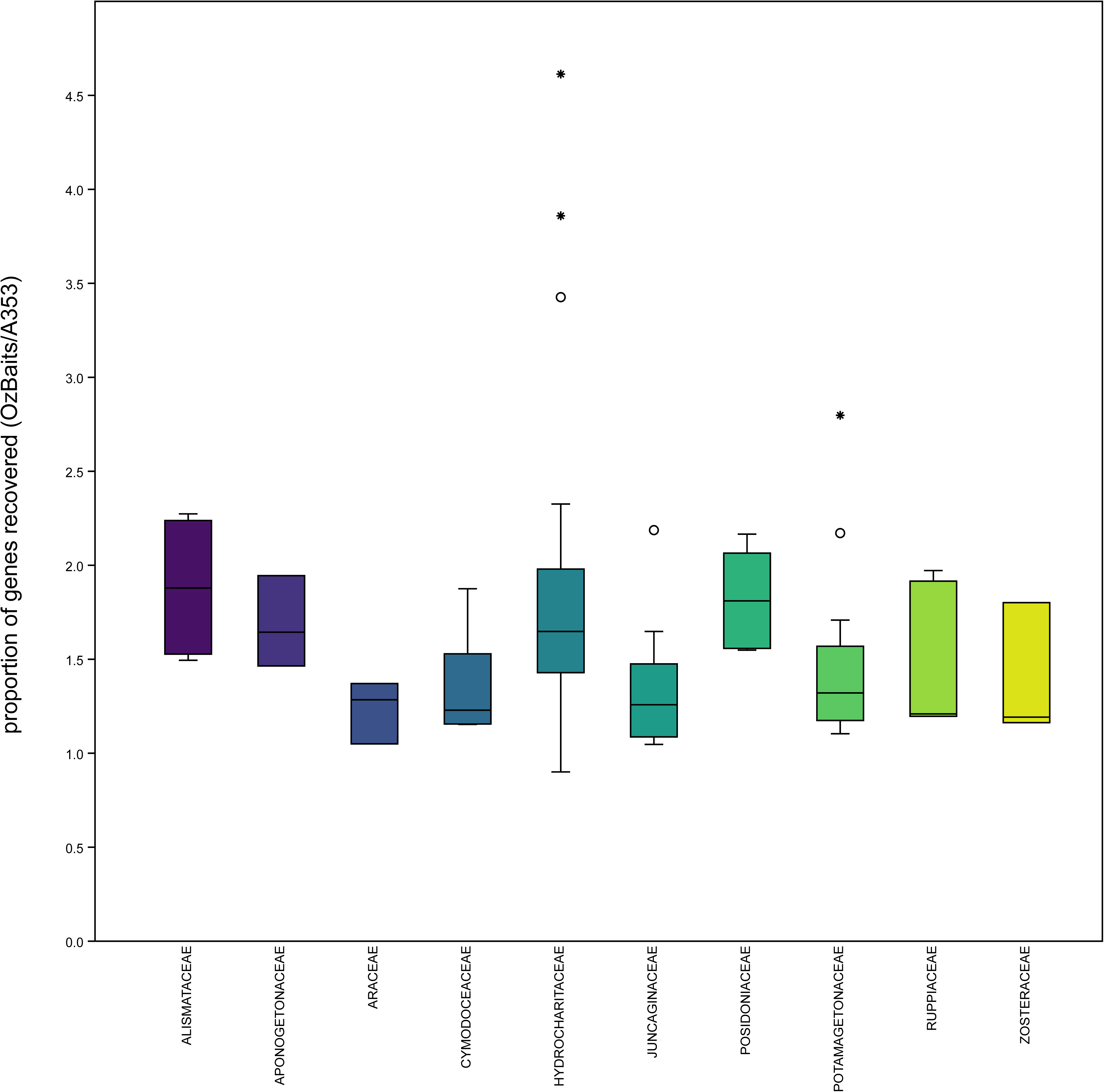
Box plots showing the relative proportion of genes recovered per sample for OzBaits and A353, ordered by family. Shown are the median (horizontal line), 25-75% quartiles (box), maximal and minimal values (whiskers) and outliers (open circle, values 1.5X IQR; asterisk, values 3X IQR).

We assessed read depth across targeted loci - for the OzBaits data the median read depth was 120.7 (*n*=98; interquartile range, IQR=81.3) while for the A353 the median was 35.9 (*n*=314; IQR=22.5). The Mann-Whitney U test indicates read depth was significantly higher for gene regions recovered by the OzBaits bait set (*z* =13.2; *p*<0.001). Similarly, the median read depth across samples was higher for OzBaits (*n*=91; median=94.9, IQR=155.5) when compared to the A353 bait set (*n=*91; median=23.3; IQR=27.4) and this comparison was also statistically significant (*z* =7.029; *p*<0.001) (Supplementary material-S8).

We found that, on average, the OzBaits bait set recovered a significantly higher proportion of the reference sequence in the ‘best hit’ contig(s) (Mann-Whitney U: *z=*54.935, *p*<.001). We also assessed the best hit LG50 and LG90 scores (i.e. the least number of contigs in the best hit that contain 50% or 90%, respectively, of the reference locus length). When considering the proportion of best hit contigs that exceed LG50 and LG90 thresholds, values for the OzBaits bait set were significantly higher for both metrics (LG50: χ^2^ =505.27, *p <* .00001; LG90: χ^2^ =2845.36, *p <* .00001). For both LG50 and LG90 ≥1, the average number of contigs in the best hit was higher for A353 (Table 2).

**Table 2:** Contig recovery for the Alismatales data set from the OzBaits and A353 bait sets. The LG50 and LG90 values indicate the least number of contigs in the best hit that contain 50% or 90%, respectively, of the reference locus length.

|  | OzBaits | A353 |
| --- | --- | --- |
| Percentage of target in best hit contig(s) [median (IQR)] | 96.43 (28.25) | 73.68 (40.31) |
| contigs with LG50 ≥1 | 10794 (87%) | 17694 (78%) |
| contigs with LG90 ≥1 | 7275 (59%) | 6780 (30%) |
| LG50 average number of contigs in best hit | 1.027 | 1.29 |
| LG90 average number of contigs in best hit | 1.07 | 1.39 |

### Phylogenetic usefulness

We used the subsampling approach described by Mongiardino Koch (2021; see also Mongiardino Koch and Thompson, 2021) to order genes by their phylogenetic usefulness as means of comparing between OzBaits and A353, but also exploring the utility of the three alignment formats output by Captus. With respect to the top-ranking alignments, we found the OzBaits data to be dominated by GF format (94%) with both NT and GE each comprising 3%. In contrast, the GF, GE and NT formats comprised 56% (175), 35% (109) and 9% (29) of the best ranked A353 alignments. In line with this, we found that phylogenetic usefulness for OzBaits was highest for GF (median rank = 70, IQR = 117) followed by GE (median = 170, IQR = 128) and NT (median = 182, IQR = 133). For the A353, both GF (median = 402, IQR = 433) and GE (median = 441, IQR = 510) ranked higher, on average, than the NT alignment format (median = 547, IQR = 427). Figure 2 show the cumulative percentage of the 407 ‘best’ ranked alignment formats for OzBaits and A353. The median ranking for OzBaits was 130 (IQR = 140) while for A353, the median was 237 (IQR = 208). Approximately 38% (37 loci) of the OzBaits and 20% (63 loci) of the A353 loci fell within the ‘best 100’ genes and this association was statistically significant (χ^2^ = 11.33, *p =* 0.00076). While the A353 data included some of the top ranked genes, approximately 60% had a rank of 200 or lower and approximately 1/3 (34%) had a rank below 300. In comparison, the OzBaits data included approximately 25% of alignments ranking below 200 (χ^2^ = 30.627, *p <* 0.00001) and just 2% of alignments below 300 (χ^2^ = 37.989, *p <* 0.00001) (Figure 2).

**Figure 2:**
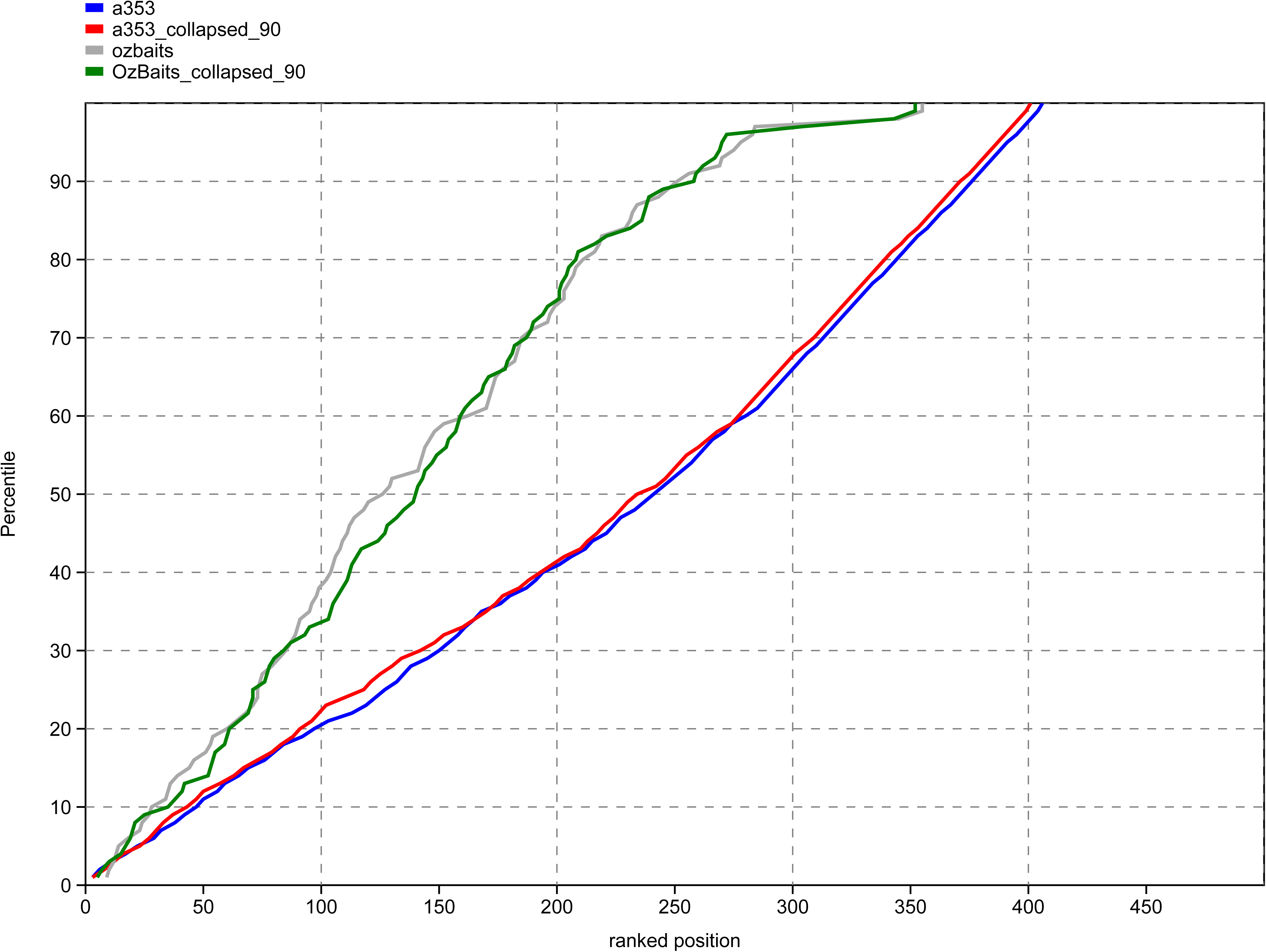
Percentile plots showing the percentage of genes with a phylogenetic usefulness ranked value lower than *x,* for ‘best’ OzBaits and A353 data sets, along with rankings for each data set estimated using a species tree with poorly supported nodes collapsed (UFBS < 90%).

We conducted three additional analyses using *genesortR* to assess the impact of (1), species tree topology (poorly supported branches collapsed in the species tree), (2) poorly aligned regions and spurious sequences (alignments cleaned using ParaGone) and (3), spurious sequences only (alignments processed using TreeShrink). With respect to (1), we found no substantive difference in the ranking of genes by phylogenetic usefulness relative to the fully resolved species tree (Figure 2). For (2) and (3), based upon Mann-Whitney U tests, we found no significant difference in phylogenetic usefulness rankings for comparisons between alignments output by Captus versus the cleaned alignments for each of OzBaits and A353, although the differences in ranking between each bait set were significant for each treatment (Supplementary material S9; Figures S4 and S5).

Table 3 shows the distribution of the 6 gene properties that were used to derive a phylogenetic usefulness axis along with an additional 6 gene properties estimated as part of the *genesortR* routine. With respect to the former, the OzBaits data show high values for the 3 proxies for signal (proportion of variable sites, average bootstrap and RF similarity) and these values are significantly different from the equivalent measures for the A353 data. In contrast, the A353 data show lower, and significantly different values for 2 proxies of bias *viz.* saturation and patristic distance. The difference in root to tip variation, the third proxy for bias, was not statistically significant for the OzBaits-A353 comparison. Across all 6 gene properties, we found that A353 dataset is characterised by high variability relative to the OzBaits and ‘best 100’ datasets.

**Table 3:**
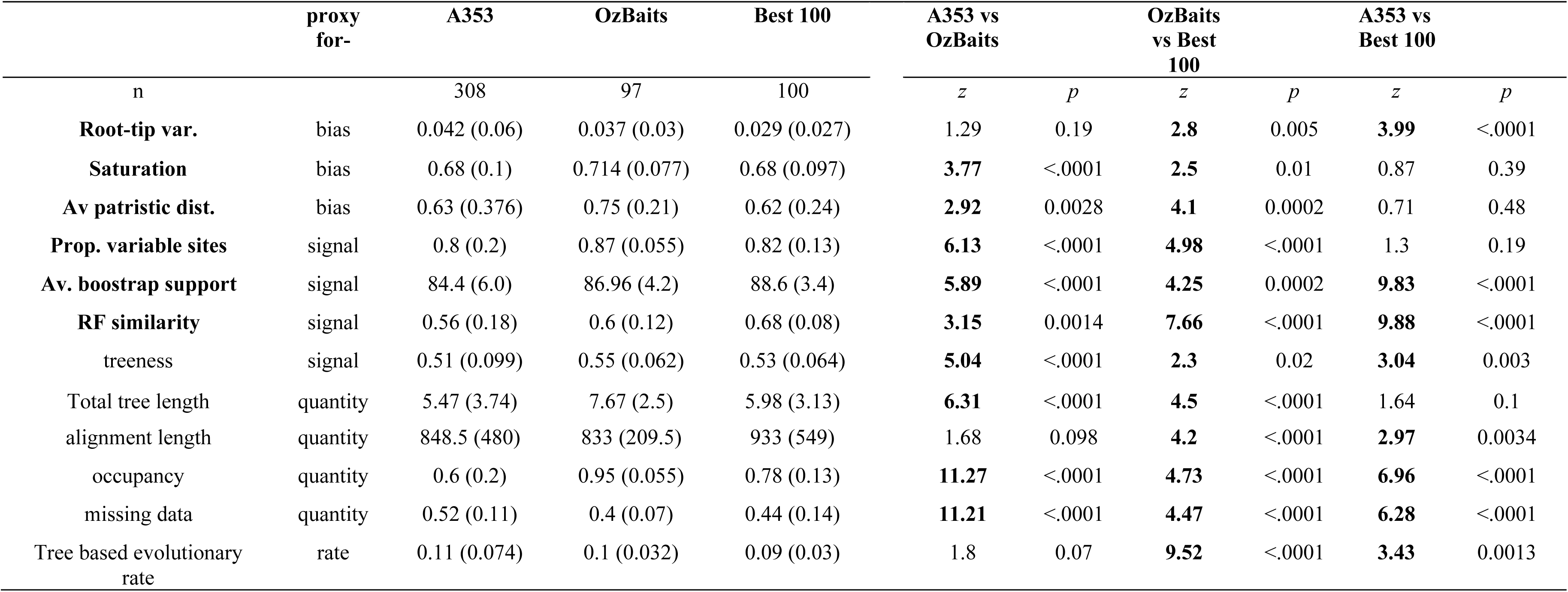
Gene properties estimated using the *GenesortR* script for the A353, OzBaits and ‘best 100’ data sets. Gene properties in bold typeface are used for ranking genes along an axis of phylogenetic usefulness, while properties that are estimated but not used for ranking are also included. Values are median (interquartile range) estimate from *n* genes included in each data set. The Mann-Whitney U test was used to assess the significance of differences between gene properties for each pairwise data set comparison. For each comparison, the *z* score and its corresponding *p* value are indicated. A *z* score > 1.96 and *p* values >0.05 are taken to indicate a significant difference (bold).

As suggested by the LG50 and LG90 statistics, above, the proportion of missing data (i.e. the proportion of missing/ambiguous cells in alignments [Mongiardino Koch, 2021]) was significantly higher for the A353 dataset relative to OzBaits and the ‘best 100’ alignments (Table 3) and shows a significant negative correlation with mean read depth (Spearmans correlation: *rs*(403) =.389, *p*<0.001). In turn, we found that missingness shows a significant negative correlation with measures of signal including RF similarity (*rs*(403) =.369, *p* < 0.001) and average bootstrap support of gene trees (*rs*(403) =.166, *p* < 0.001) and positive correlations with measures of bias including variance of root-to-tip distances (*rs*(403) =.337, *p*<0.001) and mean patristic distance (*rs*(403) =.298, *p* < 0.001).

### Phylogenetic relationships of Alismatales

We constructed 3 alignments for a common set of 85 taxa comprising ‘OzBaits best’, ‘A353 best’ and ‘best 100’, with a concatenated alignment length of 78,244, 299,665 and 116,204 bp, respectively (Supplementary material S10). Across all three data sets, phylogenetic estimates were by-and-large well-supported and congruent particularly as they relate to relationships among families and higher clades, irrespective of the inference method used. For the OzBaits data, the position of *Maundia* is incongruent relative to both the A353 and ‘best 100’ phylogenies, but with weak statistical support. Other areas of disagreement are largely confined to intra-familial and infra-generic relationships and are mostly associated with short internal branch lengths (Figure 3, Figures S1-S3).

**Figure 3:**
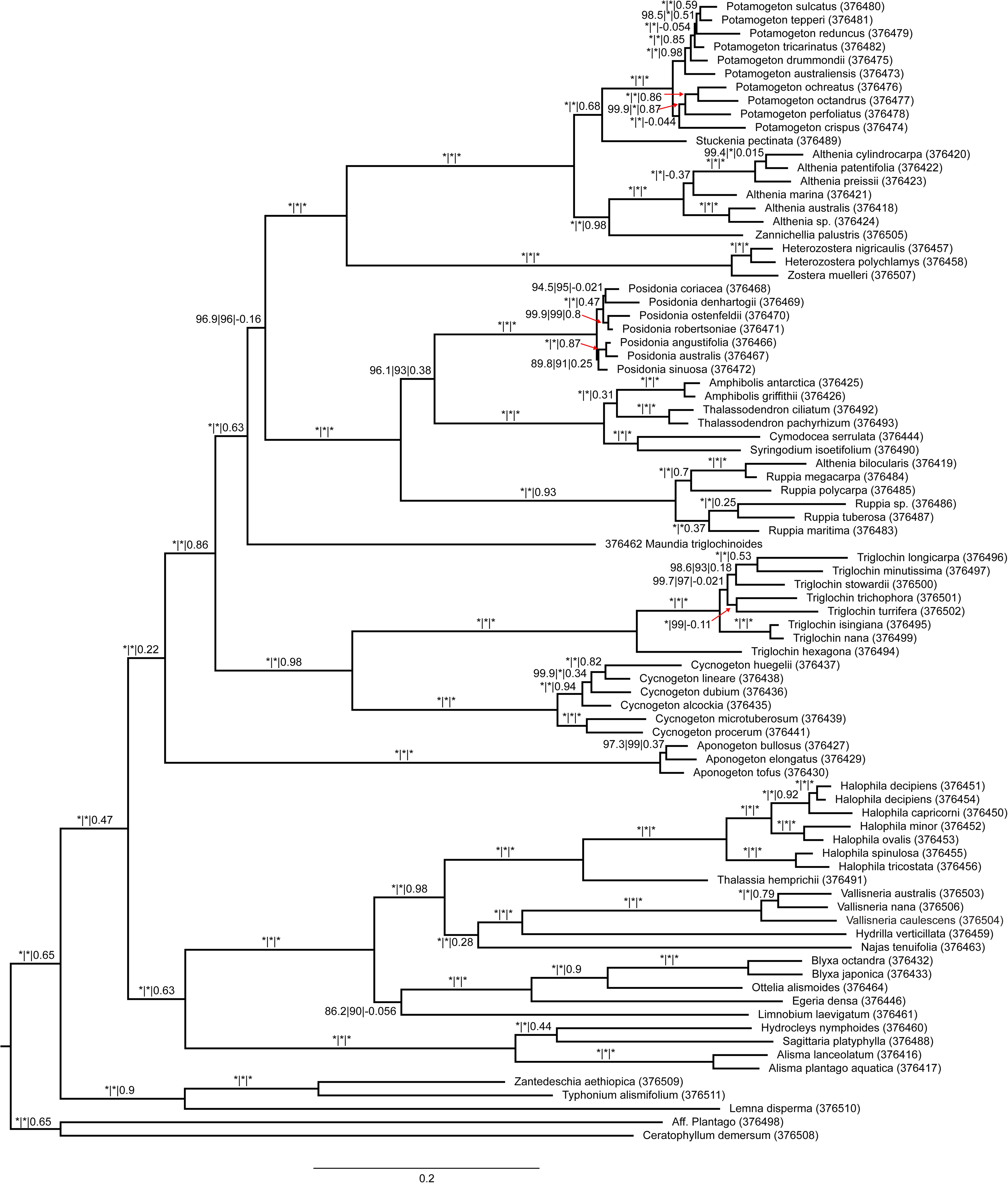
Maximum likelihood phylogeny for Alismatales and outgroups estimated from the ‘best’ 100 locus alignments, ranked according to phylogenetic usefulness, from both the OzBaits and A353 data sets. Values on the branches are support values estimated using the approximate likelihood ratio test, ultrafast bootstrap (both estimated using IQ-TREE) and quartet concordance (estimated using Quartet Sampling). Values of 100/100/1.0, respectively, are indicated by an asterisk. The GAP sample ID is indicated in brackets following the taxon label.

Quartet sampling was used to explore patterns of support and conflict in the data. As with the ML trees, evidence for discordance is largely associated with relationships within genera and above family level. With respect to the former, relationships within densely sampled lineages such as *Potamogeton*, *Posidonia*, *Triglochin* and *Cycnogeton* show high levels of discordance. Several backbone relationships, including the position of *Maundia*, are also discordant. In general, QC scores, averaged across all internal nodes, increase from A353 (0.65) to OzBaits (0.71) to the ‘best 100’ dataset (0.79) (Figure 3, Figure S1).

## Discussion

A number of ‘universal’ bait kits have been designed to target low-copy nuclear genes among vascular plants including flagellate land plants (GoFlag; Breinholt, 2021), ferns (Wolf et al. 2018), conifers (REMcon; Khan et al. 2024) and angiosperms (e.g. Buddenhagen et al. 2016; Johnson et al. 2019, A353). Of these, A353 provides a valuable comparison with OzBaits v.2, given the taxonomic scope of interest, they target a common set of genes and use the much of the same genomic resources in design. However, these two differ in the approach for selecting representative sequences for probe design, including the use of different clustering algorithms and thresholds and the proportion of a gene (transcript) targeted for each low copy locus. In the present study, we have generated data sets using OzBaits and A353 bait sets from a common set of samples, employed consistent library preparation and sequencing steps and used a common phylogenomics pipeline. By removing these confounding factors our comparisons can offer insight into how factors relating to bait design might impact on the generation of phylogenomic data using universal probe kits.

### Gene recovery: OzBaits versus A353

While the application of the A353 bait kit across angiosperms is well-established (Zuntini et al. 2024), we explored, in a limited sense, the consistency of gene recovery using the OzBaits bait set. For the angiosperm-wide assessment, gene recovery was high and extremely consistent across lineages spanning much of the extant diversity within angiosperms (Supplementary material S3) suggesting that the OzBaits target genes should be efficiently recovered from any flowering plant lineage.

In a more focused assessment, we compared locus recovery for the two baits sets across the angiosperm order Alismatales, with an estimated crown group age extending to the lower Cretaceous (Chen et al. 2022). Differences in recovery success (i.e. the proportion of genes recovered relative to the number of genes in each baits set; Fig 1, Supplementary material S7) could have several underlying causes including *in vitro* (i.e. the sequence divergence between baits and target sequences) or *in vivo* (e.g. the distance between reference and target sequences during bioinformatics processing) explanations (McLay et al. 2021). We suggest the former is more likely given that for taxa with the lowest relative recovery success from A353 (e.g. Alismataceae, Hydrocharitaceae, Posidoniaceae; Figure 1), family specific reference sequences were included in the bioinformatics processing steps (Supplementary material S2). This is further supported by significant differences in average read depth between the bait sets (Supplementary material S8)– read depth (capture efficiency) is predicted to decrease as divergence increases due to lower probe specificity (Grover et al. 2012; Portik et al. 2016; Hutter et al. 2022). According to Liu et al. (2019), the capture efficiency of low-copy nuclear genes in mosses declined sharply when the average pair-wise distance between probe and target sequences fell below 30%. For the A353 baits set, Johnson et al. (2019) use a 30% threshold of pair-wise sequence divergence in their probe design and at this value, 95% of the Angiosperm diversity was captured by 15 or less representative sequences per gene. In contrast, the OzBaits target sequences were clustered at 85% sequence similarity, which results in a higher diversity of probe sequences and presumably, largely accounts for the high consistency of target gene recovery across taxa (Supplementary material S3 and S4). On the one hand, the A353 bait set recovered an average of more than 200 genes per sample for Alismatales but uses approximately half the number of probes (75,151 120-mer probes; Johnson et al. 2019) relative to OzBaits and is therefore a highly efficient approach. On the other hand, the OzBaits probes recover fewer and (on average) shorter loci, but also have higher matrix occupancy and fewer missing values (Table 3).

### Phylogenetic usefulness: OzBaits versus A353

We used the approach developed by Mongiardino Koch (2021) to explore the properties of our data and to rank genes by their phylogenetic usefulness. While the best performing genes overall were captured by A353, a large proportion of A353 alignments also ranked poorly, including virtually all the lowest ranked 100 genes (Figure 2). We found that on average, loci recovered by the OzBaits bait set had higher phylogenetic usefulness than A353. Furthermore, a significantly higher proportion of the OzBaits targets were represented in the 100 top ranking genes relative to A353. These patterns hold for genes that were recovered by both bait sets: of the 27 recovered genes in common, c. 63% of those recovered using the OzBaits bait set ranked higher than the equivalent A353 locus (Supplementary material-S11) suggesting that the contrasting patterns of usefulness are not solely a consequence of target gene selection (e.g. selecting genes from different functional groups).

In addition to lower overall recovery of sequences per gene (i.e. matrix occupancy) (Table 3), the A353 genes have a significantly higher proportion of missing data (i.e. the individual is present in the alignment but is represented by a fragmentary sequence, type 2 missing data *sensu* Hosner et al. 2016). This is indicated by a lower proportion of the targeted regions recovered for the A353 data, and of these, fewer sequences exceed LG50 and LG90 (i.e. the best hit contig(s) exceed 50% or 90% of the target length, respectively) thresholds (Table 2). Furthermore, the A353 LG50 and LG90 best hits are more likely to include multiple contigs relative to the OzBaits data. In previous studies, it has been found that gene tree accuracy can be negatively impacted by fragmentary sequences (Hosner et al. 2016; Sayyari et al. 2017; Smith et al. 2020). Here, we found that the proportion of missing data shows significant correlations with gene tree accuracy including measures of signal (a negative correlation with RF similarity and average bootstrap support) and bias (a positive correlation with variance of root-to-tip distances and mean patristic distance) (Table 3). Fragmentary sequences can behave as ‘rogue’ taxa in gene tree estimates, and this has been attributed to lower signal because of fewer informative sites in short contigs relative to full length sequences (Goloboff and Szumik, 2015; Hosner et al. 2016).

We found that read depth is a significant correlate with the proportion of missing data in alignments although factors relating the strategy used to select the target sequences used for probe design are also likely to affect the patterns of missingness in our data. For the OzBaits bait set, the target sequences comprise a single (or partial) exon per targeted gene region, while the A353 bait set uses probes that are tiled across an entire transcript (Johnson et al. 2019). In the first instance, the recovered locus includes the targeted region and generally the upstream and downstream flanking regions comprising introns, UTRs and/or additional exon sequence. In contrast, the strategy adopted for the A353 bait set is to recover full genes including exons as well as introns/UTRs to generate ‘supercontig’ sequences (Johnson et al. 2016). One implication of the above is that, while patterns of missingness should be reasonably predictable for the OzBaits loci (i.e. a ‘core’ region corresponding to the target sequence, with the proportion of missing data increasing towards the flanking regions, see for example Streicher et al. 2016), this will be less the case for the A353 alignments. For the latter, patterns of missingness will be heterogeneous given that the majority of best hit contigs recovered a fraction of the target sequence, there was more likely to be multiple contigs in the best hit, and the target sequences commonly span multiple exons. One possible scenario is that patterns of missingness generate biased gene trees – for example, alignment portions uniquely shared by a set of distantly related terminals could artifactually inflate shared signal and lead to erroneous phylogenetic inference (Smith et al. 2020, 2023; Uribe et al. 2022). Conversely, closely related terminals could share no unique site patterns due to low alignment overlap.

Taken together, our results suggest increasing proportions missing data in alignments leads to decreased gene tree accuracy and increased bias and heterogeneity in these factors largely drive the ranking of genes by phylogenetic usefulness. While ranking using RF similarity suffers potential circularity (Mongiardino Koch, 2021), we found that species trees recovered from each of our data sets show broadly similar topologies (Figure 3 and Figures S1-S3) with respect to each other and to the target tree and poorly supported branches (largely associated with congeneric species groups) are generally held in common. Furthermore, collapsing poorly supported branches in the target tree did not substantially influence the ranking of genes (Figure 2; Supplementary material S9). In addition, we found that gene rankings correlate with alternative measures of signal (average bootstrap support) that are independent of the species tree estimate. For the OzBaits data, higher average bootstrap support relative to A353 reflects the fact that the ‘best’ loci from the former data set are overwhelmingly ‘genes-flanked’ alignments, which supports the efficiency of this bait set in capturing high coverage and largely complete contigs. In contrast, unpredictable patterns of missing data recovered by the A353 bait set appear to drive variation in gene properties and as a consequence, large variation phylogenetic usefulness.

We note that to date, the processing of A353 sequencing data has predominantly used the HybPiper pipeline (Johnson et al. 2016) and by comparison, Captus is recently developed and has yet to be fully evaluated. However, a in a recent study using sequence data simulated on the *Arabidopsis thaliana* genome, it was found that Captus recovered substantially more loci than HybPiper at low read depths (sequencing depth<10X). On the other hand, at very low sequencing depths (<5X) Captus recovered a higher proportion of shorter and less accurate contig sequences relative to higher sequencing depths (Raza et al. 2023). The recovery of short and inaccurate sequences as a consequence of low capture efficiency could potentially contribute to alignment and gene tree inaccuracy in our present data set. In order to explore this possibility, we ran the *genesortR* script on alignments that had been trimmed using the ParaGone pipeline, and separately, using TreeShrink to remove unexpectedly long branched terminals. In the case of the former, the ‘cleaned’ alignments showed lower bias and improved RF similarity but lower average values for measures of signal (proportion of variable sites and average bootstrap support). Similarly, using TreeShrink to refine alignments reduced bias but also signal. In general, we found no significant change in phylogenetic usefulness rankings for either the OzBaits or A353 data sets, although we found a small (but not statistically significant) decrease in average phylogenetic usefulness ranking for the OzBaits genes following both treatments (Supplementary material S9). Thus, while alignment trimming appears to reduce bias and gene tree estimation error (the objective of the ParaGone workflow is to generate high-quality gene trees for paralogy resolution) this comes at the expense of signal and in terms of the phylogenetic usefulness of genes, improvements in bias are presumably countered by lower signal. As both data sets were more-or-less equally impacted, we suspect that issues relating to the Captus pipeline and low read-depth sequencing data have little bearing on our main findings.

### Phylogenetic relationships of Alismatales

We found that, by-and-large, species tree estimates for Alismatales were congruent across each of our data sets irrespective of the method of inference (i.e. concatenation versus a coalescent approach) with areas of disagreement largely restricted to a few poorly supported branches (e.g. the placement of *Maundia triglochinoides* in the OzBaits tree) (Figure 3; Figures S1 and S2). While near complete support from bootstrapping or marginal posterior probabilities is expected from large scale data sets such as ours (Thompson and Brown, 2022) we found that on average, QC increased from A353 to OzBaits to our ‘best 100’ data set. Both biological and technical issues could potentially drive discordance amongst quartet topologies (Lanfear and Hahn, 2024) although here, the level of agreement between concatenation and coalescent analyses suggests that incomplete lineage sorting (ILS) is not the key factor (Hosner et al. 2016; Mirarab et al. 2016). Rather, we suggest that higher discordance on the A353 species tree topology largely mirrors the variation in gene properties noted above, with a high proportion of genes ranking poorly in terms of phylogenetic usefulness. More generally, these findings support the value of subsampling phylogenomic data given that our ‘best 100’ dataset produced a similarly well-resolved species tree relative to the more gene rich A353 data but shows lower internal discordance.

Among recent phylogenetic analyses for Alismatales, our study is perhaps most readily comparable to that of Chen et al. (2022) who use a nuclear data set comprising c. 1000 low copy nuclear orthologs to develop a hypothesis of relationships. To the extent that our sampling overlaps (Chen et al. have a high representation of Northern Hemisphere lineages versus our Australian focus) we recovered relationships amongst families and genera that are generally congruent on our ‘best 100’ tree (compare our Figure 3 and Figure 1 of Chen et al.).

## Conclusion

While there are several comparisons of hybrid-capture baits sets in the literature (e.g. ‘universal’ versus ‘taxon specific’, Larridon et al. 2020; Shah et al. 2021; Yardeni et al. 2022), we are not aware of many that have compared the design of ‘universal’ baits and how this impacts the generation of phylogenomic data (but see, for example, Hutter et al. 2022). Here, we formally introduce OzBaits v. 2, and contrast this with the widely used A353 panel. Both bait sets target angiosperms but adopt different approaches to bait design - the approach adopted for A353 enables the user to recover more loci and potentially full genes at a lower cost per base relative to OzBaits. On the other hand, we found that OzBaits can recover high quality data across a broad range of angiosperms and these data have high phylogenetic usefulness for Alismatales. The present study has focussed upon a single, albeit a highly diverse order of angiosperms. In recent studies (e.g. Zuntini et al. 2024) Alismatales is placed on a long branch as sister to the remaining monocots (excluding Acorales) and may therefore be considered a phylogenetically isolated taxon. In such specific cases, it has been found that the A353 bait set shows low capture efficiency for some genes likely because the distance to the probe sequences is too great (Johnson et al. 2019).

More generally, the utility of the A353 bait set is well-established and is not called into question here. Rather, the results of our study provide insights into possible pathways by which to increase the quality of data obtained using universal hybrid capture bait sets. Cost is a key consideration in the design of a bait set, and this is largely contingent on the number of probes required to fulfill that design. This, in turn, depends upon two (not mutually exclusive) factors: the total size (in base pairs) of the regions that are targeted, and the distance between the probes and the target DNA. Increasing the number of targeted base pairs while maintaining cost requires reducing the specificity of probes. This may come with diminishing returns as we found here for the A353 data, which included a substantial proportion of low read depth and missing data. Related to this, reducing the number of base pairs targeted could be achieved by enriching for shorter loci, such as a single exon per gene, the approach adopted for OzBaits. Significantly, we found that proxies for signal were in fact higher for the OzBaits Alismatales data (Table 3), reflecting a combination of relatively high enrichment efficiency and the consistent recovery of a high proportion of the target region including flanking introns/UTRs. More studies are needed to determine whether these issues are relevant for ‘core’ angiosperm lineages that are better represented in the A353 baits design where we would expect higher capture efficiency.

Irrespective of the above, the A353 data produced a credible estimate of Alismatales phylogeny indicating that sufficient high-quality data was obtained. However, the Alismatales includes a number of well-separated lineages that are characterised by long internal branches, a pattern that may be less challenging compared to rapid radiations where incomplete lineage sorting, gene flow and gene tree estimation errors can be conflated (Whitfield and Lockhart, 2007; Cai et al. 2021; Morales-Briones et al. 2021). The OzBaits trees showed an uncertain placement for *Maundia,* presumably reflecting insufficient signal and/or conflict in these data. However, we found that internal discordance decreased from A353 to OzBaits to our ‘best 100’ data set, suggesting that for the latter, incongruence will more likely reflect biological processes rather than gene tree estimation error. As modern phylogenomic studies seek to tackle large scale data spanning a range of divergences we suggest a trade-off between more data, and more accurate data will become increasingly relevant.

## Data availability statement

Sequence data are made available via the Bioplatforms Australia web portal (https://data.bioplatforms.com/). Associated sequence IDs are listed in Table 1.

## Supplementary Material Captions

**Supplementary material S1**: List of nuclear gene targets included in the OzBaits v2. bait set. Gene names and descriptions are based upon the *Arabidopsis thaliana* ortholog. The targeted exon (*A. thaliana* gene model), mean target length and the number of unique sequences included in the design are indicated.

**Supplementary material S2**: Reference target files used to extract target genes from each data set with the CAPTUS ‘extract’ function.

**Supplementary material S3-S6**: CAPTUS gene extraction heat maps and statistics for OzBaits ‘universal’ (S3), OzBaits Alismatales (S4) and A353 using Alismatales specific references (S5) and CAPTUS inbuilt mega353 (S6) references.

**Supplementary material S7**: Gene recovery for the Alismatales data using the OzBaits and A353 bait sets (see Figure 1).

**Supplementary material S8**: Average read depth (coverage) estimated by mapping cleaned reads back to target contigs for each sample and gene recovered by CAPTUS for the OzBaits and A353 data sets.

**Supplementary material S9**: Phylogenetic usefulness rankings (median[IQR]) for alignments output by CAPTUS using a fully resolved species tree; the same alignments following alignment cleaning using the ParaGone routine; and using Treeshrink to remove long terminal branches. Statistical tests using the Mann-Whitney U compare the CAPTUS output with each of the ParaGone and Treeshrink data sets separately for each of OzBaits and A353, as well as OzBaits versus A353 for the ParaGone and Treeshrink data. For each analysis, the input files (alignment, partition file, gene tree file and species tree) for *GenesortR* are included.

**Supplementary material S10**: Alignment and partition files used for phylogenetic inference for OzBaits ‘best’, A353 ‘best’ and 100 ‘best’ datasets.

**Supplementary material S11**: Phylogenetic usefulness rankings for genes that are shared between OzBaits and A353 bait sets. For each gene, the A353 and OzBaits identifier are used in that order, and the higher ranked version is highlighted with bold typeface. Values for the 6 properties used to derive a phylogenetic usefulness axis are also included.

**Figure S1**: Maximum-likelihood topologies (IQTREE) inferred from concatenated alignments for (A) OzBaits best, and (B) A353 best data sets. Values on the branches are support values estimated using the approximate likelihood ratio test, ultrafast bootstrap (both estimated using IQ-TREE) and quartet concordance (estimated using Quartet Sampling). Values of 100/100/1.0, respectively, are indicated by an asterisk. The GAP sample ID is indicated in brackets following the taxon label.

**Figure S2**: Species tree topologies inferred using ASTRAL for (A) OzBaits best, and (B) A353 best data sets. Values on branches are local posterior probabilities (LPP). The GAP sample ID is indicated in brackets following the taxon label.

**Figure S3**: Species tree topologies inferred using ASTRAL for the 100 best data set. Values on branches are local posterior probabilities (LPP). The GAP sample ID is indicated in brackets following the taxon label.

**Figure S4**: Percentile plots showing the percentage of genes with a phylogenetic usefulness ranked value lower than *x,* for ‘best’ OzBaits and A353 data sets, along with rankings for each data set estimated following the alignment cleaning routines implemented by the ParaGone pipeline.

**Figure S5**: Percentile plots showing the percentage of genes with a phylogenetic usefulness ranked value lower than *x,* for ‘best’ OzBaits and A353 data sets, along with rankings for each data set estimated following the removal of abnormally long branches among gene trees using TreeShrink.

