## Supplementary figures and images for "A comparison of two universal angiosperm bait sets and the phylogenomics of Alismatales"

### Supplementary Figure S1

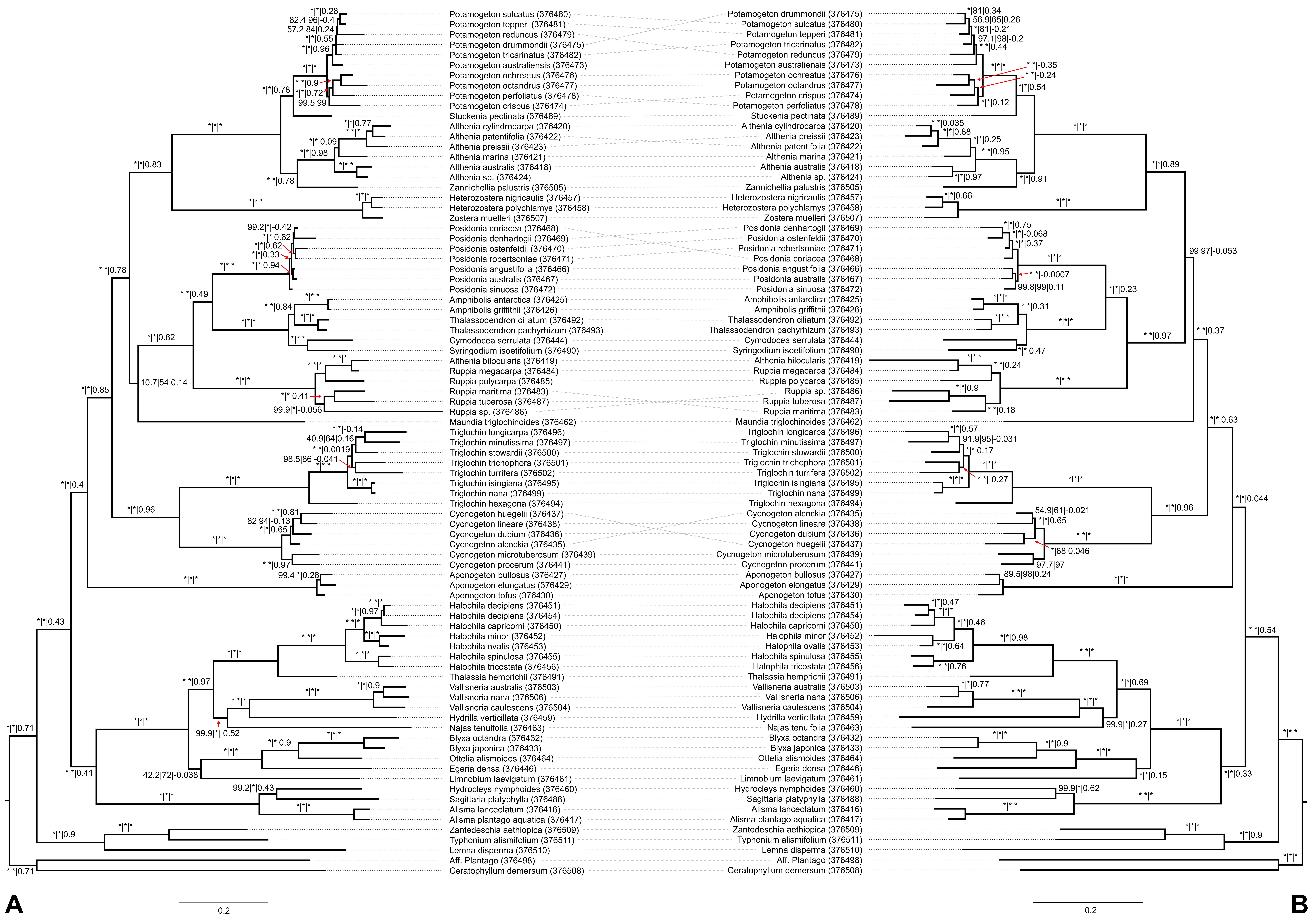

### Supplementary Figure S2

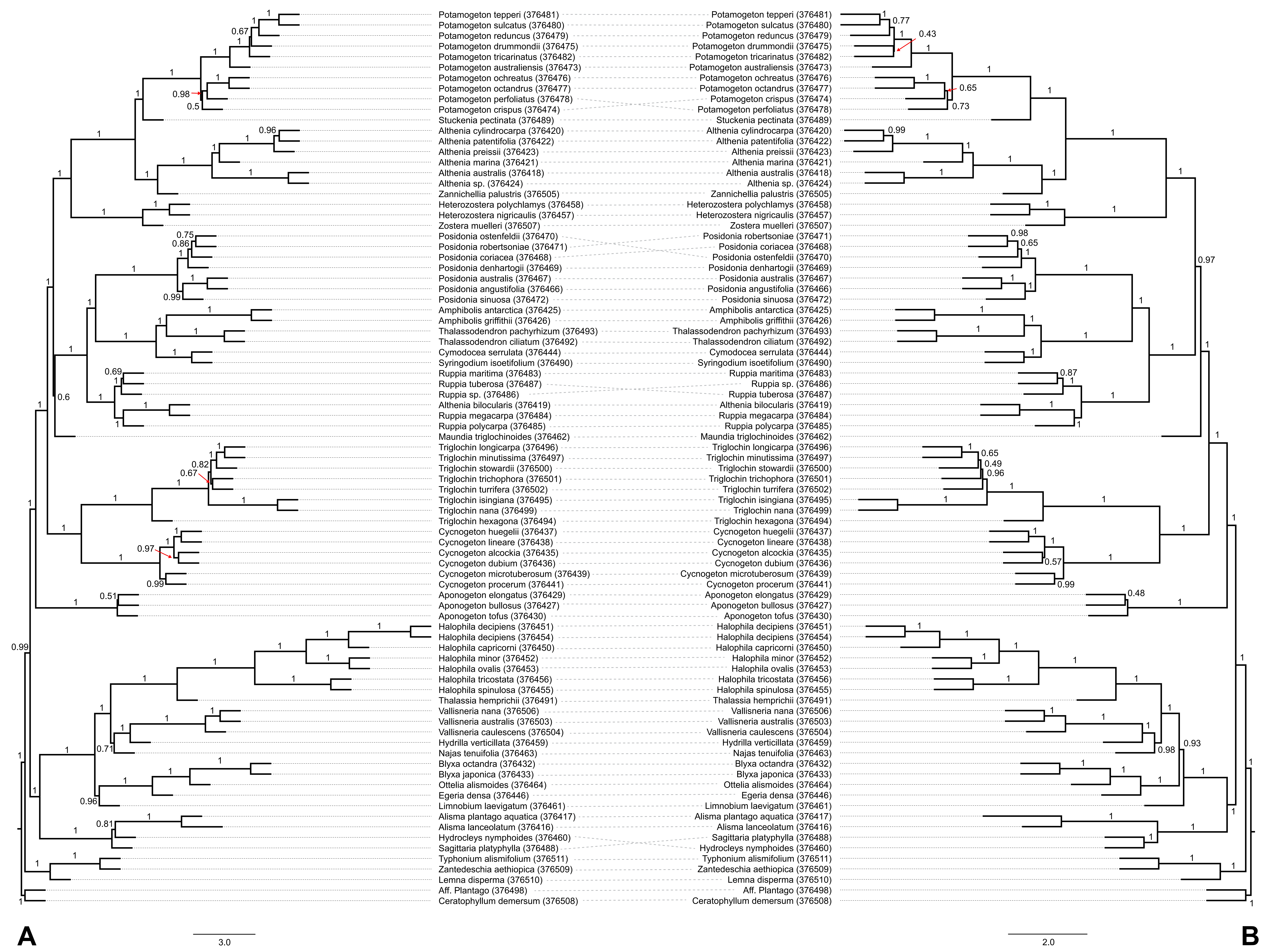

### Supplementary Figure S3

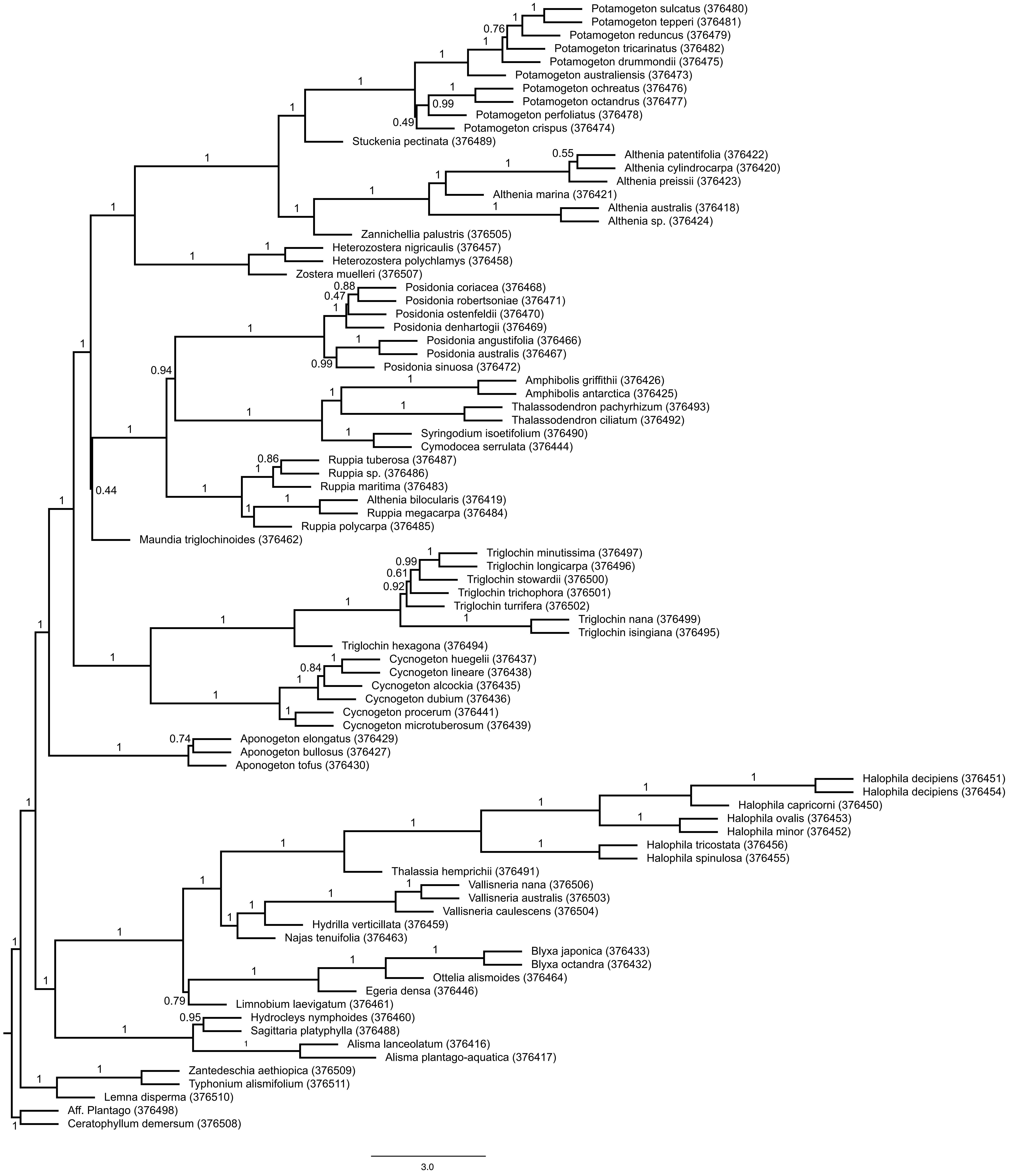

### Supplementary Figure S4

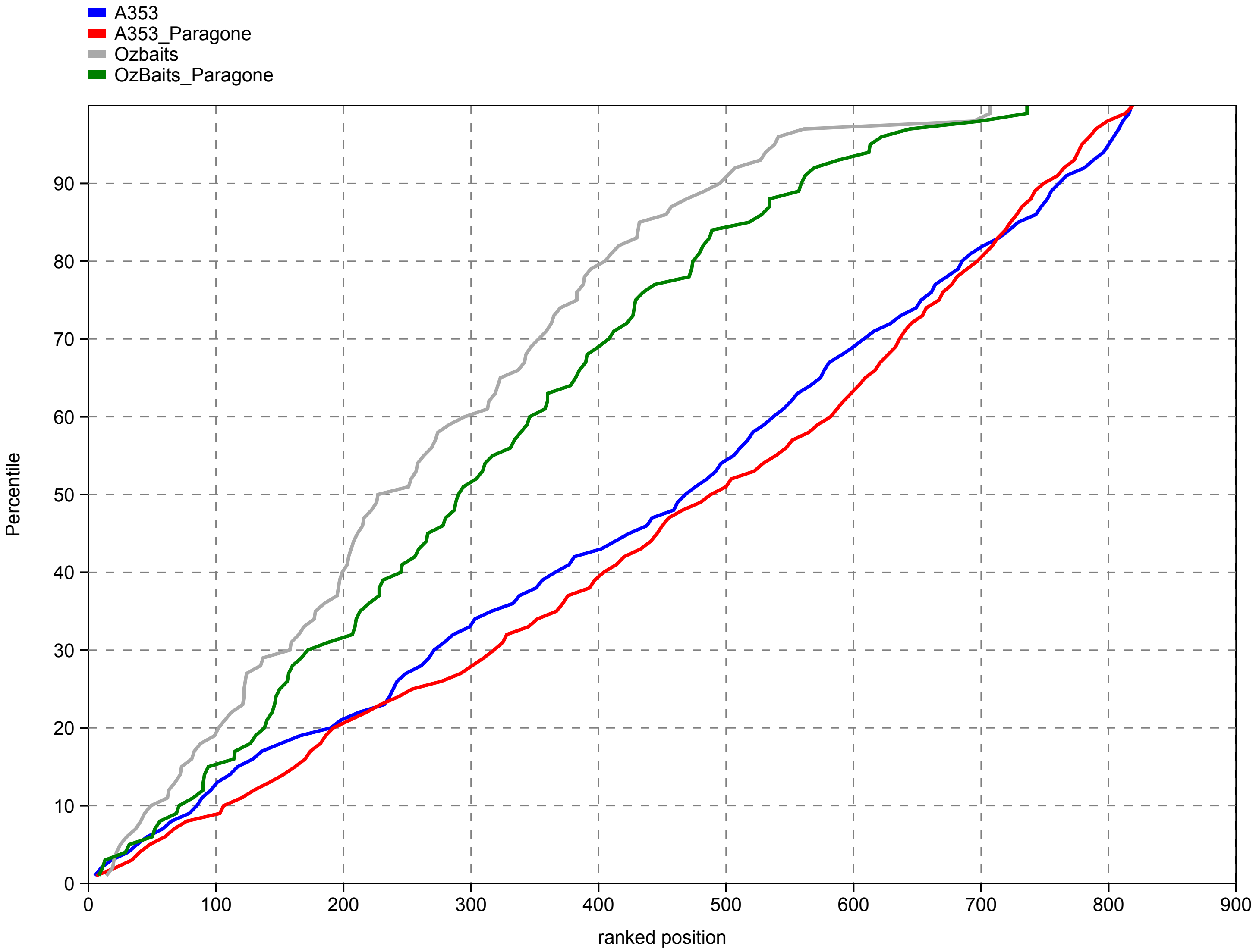

### Supplementary Figure S5

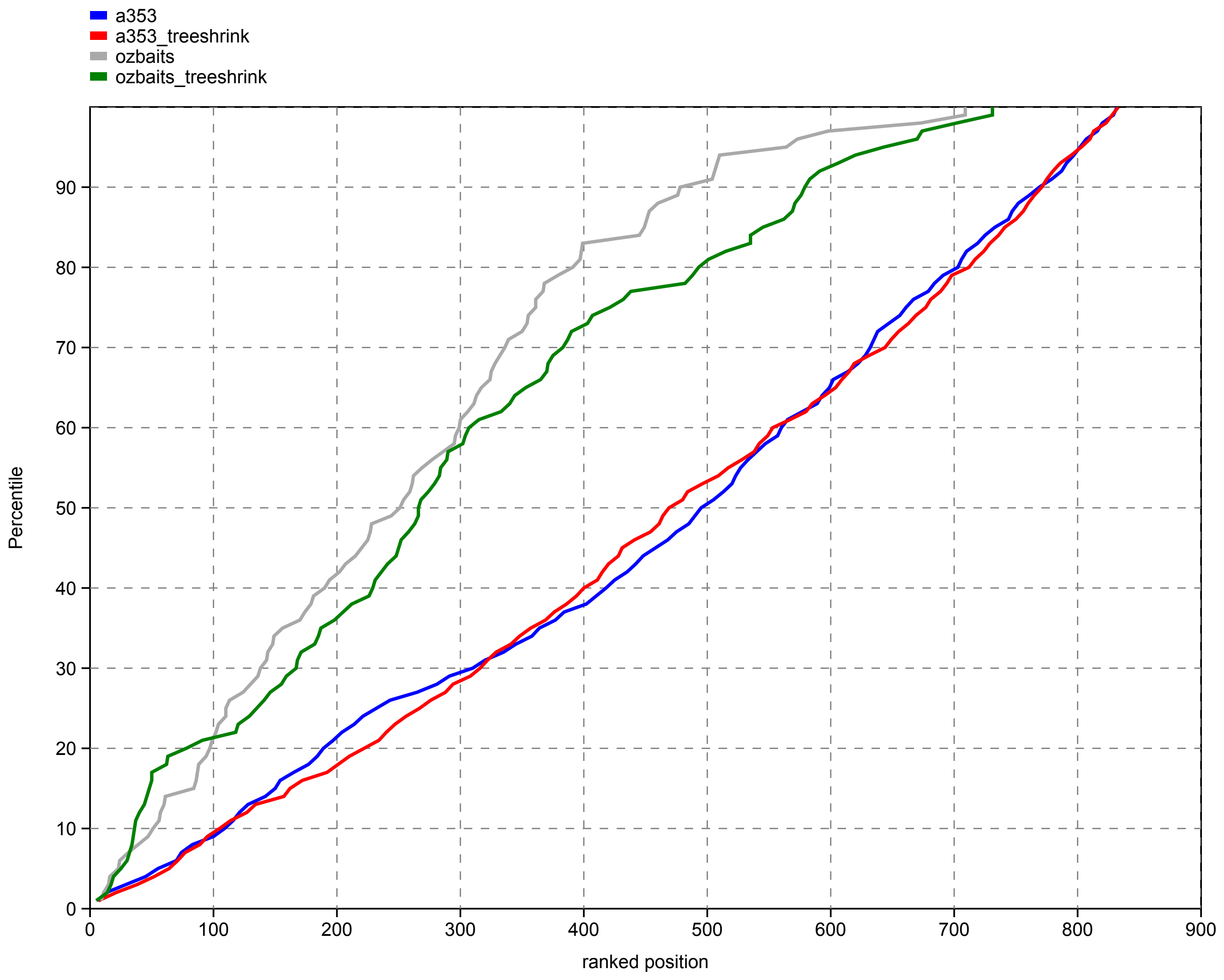
